# RNase HI depletion strongly potentiates cell killing by rifampicin in mycobacteria

**DOI:** 10.1101/2021.07.14.452003

**Authors:** Abeer Al-Zubaidi, Chen-Yi Cheung, Gregory M. Cook, George Taiaroa, Valerie Mizrahi, J. Shaun Lott, Stephanie S. Dawes

## Abstract

Multidrug resistant (MDR) tuberculosis (TB) is defined by the resistance of *Mycobacterium tuberculosis*, the causative organism, to the first-line antibiotics rifampicin and isoniazid. Mitigating or reversing resistance to these drugs offers a means of preserving and extending their use in TB treatment. R-loops are RNA/DNA hybrids that are formed in the genome during transcription, and can be lethal to the cell if not resolved. RNase HI is an enzyme that removes R-loops, and this activity is essential in *M. tuberculosis:* knockouts of *rnhC*, the gene encoding RNase HI, are non-viable. This essentiality supports it as a candidate target for the development of new antibiotics. In the model organism *Mycolicibacterium smegmatis*, RNase HI activity is provided by two RNase HI enzymes, RnhA and RnhC. We show that the partial depletion of RNase HI activity in *M. smegmatis,* by knocking out either of the genes encoding RnhA or RnhC, led to the accumulation of R-loops. The sensitivity of the knockout strains to the antibiotics moxifloxacin, streptomycin and rifampicin was increased, with sensitivity to the transcriptional inhibitor rifampicin strikingly increased by nearly 100-fold. We also show that R-loop accumulation accompanies partial transcriptional inhibition, suggesting a mechanistic basis for the synergy between RNase HI depletion and transcriptional inhibition. A model of how transcriptional inhibition can potentiate R-loop accumulation is presented. Finally, we identified four small molecules that inhibit recombinant RnhC activity and that also potentiated rifampicin activity in whole-cell assays against *M. tuberculosis,* supporting an on-target mode of action, and providing the first step in developing a new class of anti-mycobacterial drug.

**Importance:** This study validates mycobacterial RNase HI as a druggable, vulnerable candidate for a new therapeutic treatment of *M. tuberculosis* with a novel mode of action. RNase HI depletion shows synergistic bacterial killing with some current first- and second-line antibiotics, suggesting that RNase HI inhibitors would combine well with these regimens, and could potentially accelerate the clearance of drug-sensitive strains. RNase HI inhibitors also have the potential to reduce the effective dose of rifampicin, with the comcommitant reduction in side effects. The potentiation of rifampicin efficacy conferred by RNase HI deficiency suggests that RNase HI inhibitors may be able to mitigate against development of rifampicin resistance. The synergy may also be able to reverse rifampicin resistance, rescuing this antibiotic for therapy. The surprising finding that low levels of transcriptional inhibition potentiate R-loop formation provides a key new insight into R-loop metabolism.

## Introduction

The emergence of drug-resistant bacteria has been recognised as a global health crisis (1). If left unchecked, antibiotic resistance threatens to overturn the substantial gains that have been made to human healthcare by the use of antibiotics to treat and prevent infectious disease, resulting in huge social and economic costs. Globally, the antibiotic resistance crisis has been spearheaded by the alarmingly rapid rise in the prevalence of *M. tuberculosis* strains that are resistant to the cornerstone antibiotics of first-line TB therapy, rifampicin (which inhibits transcription) and isoniazid (which inhibits cell wall biosynthesis). The situation has been exacerbated by the emergence and spread of extensively drug-resistant (XDR) *M. tuberculosis* strains that are also resistant to second-line TB therapeutics, such as the fluoroquinolones (which inhibit DNA topoisomerases) and linezolid (which inhibits the ribosome). It is estimated that there were at least 450,000 new cases of MDR-TB worldwide in 2019, and at least 180,000 deaths from MDR-TB were reported (2). The World Health Organisation (WHO) has set a goal to eliminate TB by 2035, but this clearly cannot be achieved without addressing antibiotic resistance. Recent striking progress has been made towards this: in 2019, the U.S. Food and Drug Administration (FDA) approved a new combination therapy composed of bedaquiline, pretomanid, and linezolid that able to treat both MDR- and XDR-TB disease effectively using three novel antibiotics, but with higher cost and greater side effects than standard first-line therapy (3). There have been no similar efforts towards improvements in first-line therapy, which is the primary line of defense. Clearly, retaining or improving the efficacy of the inexpensive first-line drugs for the treatment of drug-sensitive TB is imperative, as is enhancing their activity where possible to reduce side effects and thereby improve patient adherence.

Genome maintenance is essential for bacterial survival. Both transcription and replication processes inflict stresses that challenge genome topology and integrity. R-loops, which are RNA/DNA hybrids that form spontaneously in the genome during transcription, are a major threat to genome stability and can be lethal if not resolved (4, 5) (Fig 1A). Persistent R-loops can alter DNA topology, block both transcription and replication, and promote double-strand DNA breaks (DSBs) (6–9). Not all R-loops are pathological, since R-loop formation can be a necessary intermediate in plasmid replication (10) and in bacterial immune system surveillance via CRISPR (Clustered Regularly Interspersed Short Palindromic Repeats) (11). Multiple helicases and endonucleases can contribute to the resolution of these hybrids, but ribonuclease HI (RNase HI) activity provides the major dedicated function for this, by targeted hydrolysis of the RNA strand in an RNA/DNA hybrid (12). Cells therefore require precisely controlled levels of RNase HI to prevent dysfunction. Recently it was demonstrated that loss of RNase HI function in *E. coli* drives the extinction of rifampicin and streptomycin-resistant strains (13), suggesting that RNase HI might be a promising new antimicrobial target in light of the current antimicrobial resistance crisis. RNase HI is already a validated target for HIV therapy: Much progress has been made over the last two decades in the discovery and optimisation of inhibitors with different binding modes that are specific and selective to the RNase H activity of HIV-1 RT and that have antiviral activity within cells (14–16).

**Figure 1.**
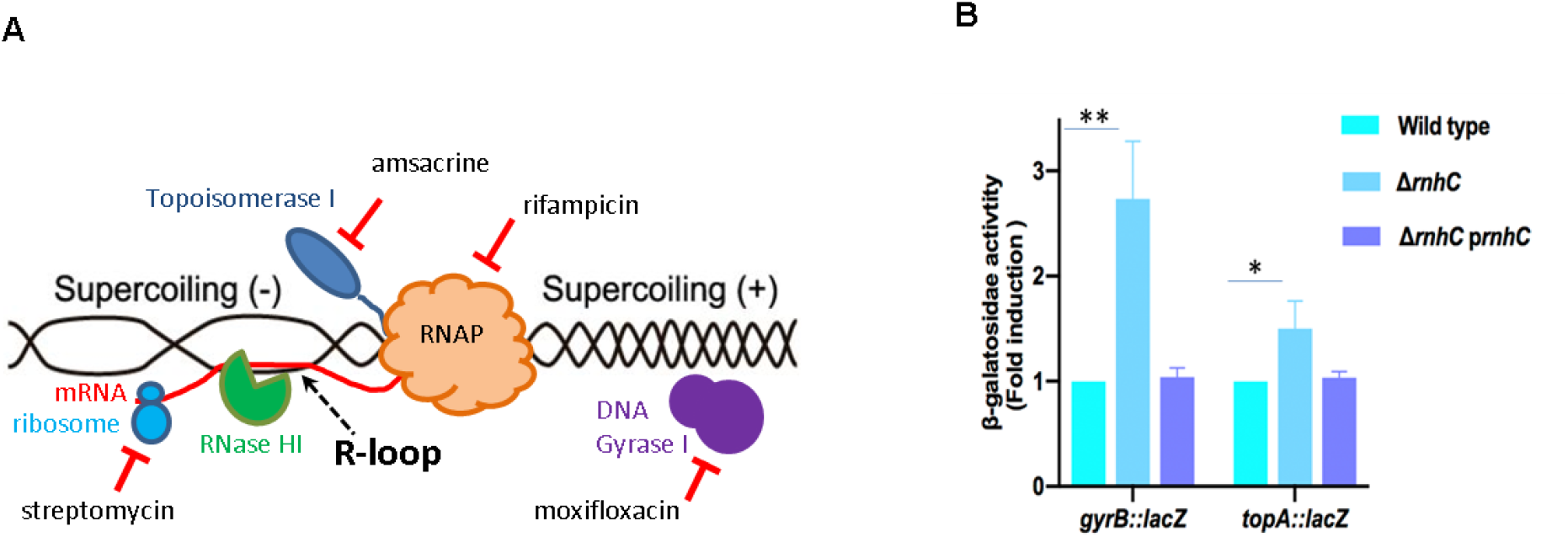
**A**. Twin domain model of transcription and location of enzymes (and their inhibitors) involved in topological modification or R-loop metabolism. RNAP: RNA polymerase I. **B**. Expression of gyrase (*gyrB*) and topoisomerase I (*topA*) promoters fused to *lacZ* in wild type, Δ*rnhC* and complemented strains of *M. smegmatis* mc^2^155. The relative activities of the β-galactosidase reporters are shown, normalized to the wild type. Data shown are representative of the average of three independent experiments with standard deviations indicated by error bars. The statistical significance of the differences assessed using Student’s unpaired *t*-test calculated by GraphPad Prism 7. \**, P* < 0.05; **, *P* < 0.01.

The mycobacteria possess a bifunctional RNase HI enzyme called RnhC that is composed of an N-terminal domain with RNase HI activity and a C-terminal domain from the acid phosphatase family. This domain possesses cobalamin-P phosphatase activity (CobC) and is proposed to be involved in vitamin B_12_ biosynthesis (17–20). These domains can function separately, although the CobC domain confers increased activity on the RNase HI domain of *M. tuberculosis* RnhC (Rv2228c) (17). The mechanism for this is not understood. Although most mycobacteria have only one gene encoding RNase HI activity, the non-pathogenic saprophytic model organism, *M. smegmatis*, also possesses a single-domain RNase HI termed RnhA in addition to RnhC. Both RNases HI have been well characterised biochemically, as have single knock-out strains of both *rnhA* and *rnhC* (18, 19, 21), which both grow indistinguishably from wild type (18, 19). However, simultaneous deletion of both genes is lethal, indicating the essentiality of RNase HI activity in mycobacteria (18, 19). Transposon mutagenesis in *M. tuberculosis* located essentiality of function to the RNase HI domain of RnhC (22) (23). Moreover, *M. tuberculosis rnhC* can rescue *M. smegmatis* from the lethality of the double knockout of *rnhA* and *rnhC* (20).

In this study we used the dual RNase HI capacity of *M. smegmatis* provided by RnhA and RnhC to investigate RNase HI as a novel drug target in mycobacteria. We used single RNase HI knock-out strains to deplete RNase HI activity, and showed that deletion of RnhC in particular is sufficient to cause the accumulation of R-loops, and to induce markers of genome topological stress. Furthermore, deletion of RnhC enhanced the killing activity of various anti-tubercular drugs, with an especially marked effect on rifampicin activity. We also identified four small molecules, previously shown to inhibit HIV RNase HI, as micromolar inhibitors of recombinant *M. tuberculosis* RnhC *in vitro*, that also potentiate rifampicin killing in whole *M. tuberculosis* cells. Together, these results validate RNase HI as a drug target in the mycobacteria, demonstrate the potential for enhanced therapy by combining rifampicin treatment with RNase HI inhibitors, and open avenues to rescue rifampicin-resistant strains for therapy with rifampicin through RNase HI inhibition.

## Materials and methods

### Strains, growth conditions and media

Strains and plasmids used in this study are listed in Tables S1 and S2. *E. coli* strains Top10 and BL21(DE3)pLysS, used for cloning and protein expression respectively, were grown in lysogeny broth (LB) and/or agar at 37°C or 28°C in the presence of the appropriate antibiotic(s) at the required concentrations *i.e.* 50 μg ml^−1^ kanamycin (Goldbio), 100 μg ml^−1^ ampicillin (Goldbio), 200 μg ml^−1^ hygromycin (Thermofisher), 5 μg ml^−1^ gentamicin (Goldbio) and 35 μg ml^−1^ chloramphenicol (Sigma-Aldrich). *M. smegmatis* mc^2^155 and *M. tuberculosis* H37Rv were grown at 37°C in Middlebrook 7H9 broth (BD) or on Middlebrook 7H10 agar (BD) supplemented with 10% Albumin-Dextrose-Saline, 0.2% glycerol and 0.05% Tween 80 (Sigma) in the presence of appropriate antibiotics at the following concentrations: 25 μg ml^−1^ kanamycin, 50 μg ml^−1^ hygromycin and 5 μg ml^−1^ gentamicin.

### Plasmid and Strain construction

Plasmids were constructed using standard cloning techniques. Inactivation of *rnh* alleles was carried out by homologous recombination with suicide plasmids (24) containing alleles inactivated by insertion of either a hygromycin antibiotic resistance cassette (Δ*rnhC*) or kanamycin resistance cassette (Δ*rnhA*). Allele replacement was confirmed initially by Southern blot and subsequently by whole-genome sequence analysis. Bioluminescent strains were constructed by transformation with the integrating plasmid pMV306G13+Lux which carries the *luxABCDE* operon (25). *M. smegmatis rnhA* or *M. tuberculosis rnhC* genes for complementation studies were cloned into the pAL5000-based replicating plasmids (26) pOLYG or p32GoriM, respectively, to make p*rnhA* and p*rnhC*. These vector backbones differ only in the antibiotic resistance cassette, hygromycin or gentamycin respectively (27). The *M. smegmatis rnhA* gene was cloned with 350 bp of upstream region containing its own promoter (28), and *M. tuberculosis rnhC* was expressed from the well-characterised leaderless Ag85 promoter (29).

### Immunodetection of RNA/DNA hybrid

Bacterial strains were grown as above and harvested at mid-log phase (OD_600_ =1.5). Aliquots (20 ml) of each culture were harvested and the nucleic acid was extracted using the phenol-chloroform technique (30) with a slight modification: Following lysozyme treatment, RNase A (10μg/ml) was added to degrade single-stranded RNA, and small RNAs were precipitated by 30% (w/v) PEG 8000/30 mM MgCl_2_. Nucleic acid was quantified using a spectrofluorometric method using QuantiFluor^®^ dsDNA (Thermofisher). Nucleic acid (400 ng) was spotted on to Amersham Hybond N+ nylon membrane (GE Healthcare Lifesciences), air-dried and UV cross-linked for 3 min using a transilluminator. Control aliquots were pre-incubated with 10 U of *E. coli* RNase H (NEB) for two hours at 37 °C before spotting. The membrane was blocked in TBST (20mM Tris pH 7.4, 150mM NaCl and 0.1% (v/v) Tween 20) containing 5% (w/v) milk powder at room temperature for 1 hour. The blocking solution was removed and replaced with fresh TBST containing 5% (w/v) milk and S9.6 anti-DNA-RNA Hybrid Antibody (Kerafast) at 1μg/ml and incubated at 4°C overnight. The following day, the membrane was washed (3 x 5 min) with fresh TBST, and finally incubated with 1:1000 dilution of HRP-conjugated anti-mouse secondary antibodies (Invitrogen) in TBST for 1 hour at room temperature. Unconjugated secondary antibody was removed by washing (3 x 5 min) with fresh TBST, and developed with West Pico PLUS Chemiluminescent Substrate (Thermofisher) according to manufacturer instructions. Chemiluminescence was detected using an Amersham Imager 600, and dot intensity was quantified using the Image Lab software suite (BioRad). Student’s unpaired *t*-test, calculated using GraphPad Prism was used to test for significance.

### β-galactosidase reporter assay

Cultures of *M. smegmatis* mc^2^155 and Δ*rnhC* were grown as above and harvested at mid-log phase. Optical density was used for normalisation of β-galactosidase activity. The assay was carried out according to Miller with variations (31, 32). Briefly, aliquots (2 ml) from each culture were harvested and resuspended in Z-buffer (60mM Na_2_HPO_4_, 40mM NaH_2_PO_4_, 10mM KCl, 1mM MgSO4, 50 mM β-mercaptoethanol) to a final volume of 600μl. Cells were permeabilized by adding 100μl of chloroform and 50 μl of 0. 1% SDS, and the mixture was incubated at 28°C for 5 minutes. Then 200μl of ONPG (4 mg/ml) was added to the mixture, which was incubated at 28°C until sufficient yellow colour developed, and time of incubation was recorded. The reaction was stopped by the addition of 500μl of 1 M Na_2_CO_3_ then the mixture was centrifuged at 10 000 g for 2 min. The OD420 of the supernatant was measured and the β-galactosidase activity was expressed in Miller Units using the following equation: OD_420_(1000)/(tv) OD _600_, where t = incubation time (min) and v = volume of culture (ml). The Student’s unpaired t-test embedded in GraphPad Prism software was used to test for significance.

### Growth inhibition assays

#### M. smegmatis

Two different methods were used to assess growth inhibition.

##### Agar proportion method

*M. smegmatis* strains were grown and harvested at mid-log phase (OD_600_ = 1 to 1.5). A single-cell suspension was prepared by centrifugation of the culture at 2,000 x g for 10 min to pellet clumps, the supernatant was collected, and 20% glycerol was added. Aliquots were frozen with liquid nitrogen and stored at −80 ⁰C. For killing assay, 150 CFU were plated in Middlebrook 7h10 agar plates containing two-fold serial dilutions of rifampicin and isoniazid and incubated at 37°C for 3 days. Bacterial survival at each concentration was assessed and MIC determination was calculated from non-linear regression analysis using GraphPad Prism. Significant differences were assessed using the Student’s unpaired t-test embedded in GraphPad Prism software.

### Bioluminescence reporter assay

Bacterial strains carrying pMV306G13+Lux were grown as above to mid-log phase (OD_600_ = 1.5), and diluted 30-fold to OD_600_ = 0.05. All assays were set up in 96 well plates (Greiner Bio-One) in a total volume of 100μL where 50μL of the diluted culture was added to 50μL of two-fold serially diluted antibiotics in Middlebrook 7H9 media. Two controls were added, bacteria only and media only. The plates were covered with lids, sealed with parafilm to prevent evaporation, and were incubated at 37°C for 18 hours. Bioluminescence was quantified using a SpectraMax iD3 Plate Reader. Bacterial survival at each concentration was calculated based on the ratio of the RLU in the presence or absence of the inhibitor. Dose response curves and the MIC50 for each drug was determined by fitting the data to log (inhibitor) vs. response – variable slope equation using GraphPad Prism 7. The Student’s unpaired t-test embedded in GraphPad Prism software was used to test for significant differences in the killing between wild type and Δ*rnhC*.

#### M. tuberculosis

*M. tuberculosis* mc^2^6230 (Δ*RD1* Δ*panCD*) (33) was grown in Middlebrook 7H9 broth supplemented with OADC (0.005 % oleic acid, 0.5 % bovine serum albumin, 0.2 % dextrose, 0.085 % catalase), 0.05 % tyloxapol and 25 μg mL^−1^ pantothenic acid (PAN) at 37 °C. The assay was carried out in 96 well plates in a total volume of 100μL. The cultures were diluted to OD_600_ = 0.05 and 50μL of the diluted culture was added to 50μL of two-fold serially diluted inhibitor in Middlebrook 7H9 media. Plates were incubated without shaking at 37 °C for 7 days before the minimal inhibitory concentrations that inhibited 90% of growth (MIC_90_) were determined from the visual presence or absence of growth in biological triplicates. The fractional inhibitory concentration index (FICI) was determined for RNase HI inhibitors in combination with rifampicin by the addition of a sub-MIC concentration of rifampicin (5 nM) to the MIC assay. FICI values were calculated using the equation FICI = FIC_A_ + FIC_B_ = (C_A_/MIC_A_) + (C_B_/MIC_B_), where MIC_A_ and MIC_B_ are the MICs of drugs A and B alone, and C_A_ and C_B_ are the concentrations of the drugs in combination.

### Recombinant RnhC purification

The *RnhC/Rv2228c* open reading frame (ORF) was amplified from *M. tuberculosis* DNA and cloned into pMAL-C2 (34) as an N-terminal maltose-binding protein (MBP) fusion construct which was further modified to include a 6xHis-tag following the MBP ORF and a 3C protease recognition sequence (35) immediately upstream of *RnhC*. The plasmid was transformed into *E. coli* BL21(DE3) pLysS (36). Cultures grown in ZYP-5052 medium (37) were auto-induced by incubation for 7 hr at 37 °C followed by incubation at 28 °C for 20 hr. The cells were harvested (5,300 x g for 30 min at 4 °C) and re-suspended in lysis buffer (20 mM HEPES and 150 mM NaCl, pH 7.5) containing 10μg/ml of DNase 1 and complete Mini EDTA-free protease inhibitor tablets (Roche) (one tablet/10ml total volume). The cells were lysed using a French press cell disruptor (ConstantSystem) at a pressure of 18 kpsi. The lysate was centrifuged at 20,000 × g for 30 min at 4 °C. Recombinant protein was purified by affinity chromatography using an amylose column, and cleaved from the tags (MBP and 6X-His) using His-tagged recombinant 3C protease (38) overnight at 4 °C. The 3C protease and the tags were removed by reverse affinity chromatography. Recombinant Mt RnhC was concentrated and further purified by size exclusion chromatography using a 16/600Superdex^®^ 200 pg columns (GE Healthcare). Mt RnhC eluted in a single peak and the fractions were pooled, concentrated to 10 mg.ml^−1^ and stored at −20°C.

### HIV-RNase HI Inhibitors

HIV RNase H inhibitors were selected from the PubChem database and kindly provided by the National Institutes of Health (NIH) through the Developmental Therapeutics Programme (DTP). The compounds were resuspended in DMSO at a final concentration of 10 mM and stored at −20°C.

### RNase HI inhibition assay

The RNase HI activity assay was performed using a Fluorescence Resonance Energy Assay (FRET) as previously described in (39) using an 18bp RNA/DNA, where the RNA strand was labelled with fluorescein on the 3′-end (5’-GAU CUG AGC CUG GGA GCU-fluorescein-3’) and the complementary DNA strand labelled with Iowa black on the 5′-end (5’ -Iowa black-AGC TCC CAG GCT CAG ATC-3’). Mt RnhC (4nM) was incubated with 25nM of substrate in FRET buffer (50 mM Tris-HCl, pH 8, 60 mM KCl, 5mM MgCl2). The increase in fluorescein signal due to substrate hydrolysis was monitored at an excitation/emission of 490nM/528nM using a SpectraMax iD3 Plate Reader. Compounds active at an initial inhibitor concentration of 100 μM were subsequently assessed using a dose response assay. Drug dose-response curves and IC50 determinations were carried out by non-linear regression analysis in GraphPad Prism 7.

## Results

### Loss of RnhC induces markers of DNA topology stress in *M. smegmatis*

As a first indicator of whether depletion of RNase HI activity affected genome maintenance, we looked at the expression of topoisomerases responsible for topology control. In metabolically active cells, R-loops can stall RNA polymerase complexes and increase topological stress in the genome because persistent R-loops prevent the resolution of both the negative DNA supercoiling that is formed in the wake of an RNA polymerase, and the positive DNA supercoiling that is generated ahead of it (40, 41) (Figure 1A). Both the DNA gyrase and topoisomerase I promoters in *M. smegmatis* respond directly to the topological stresses experienced by the chromosome since RNA polymerase binding elements in each promoter adopt optimal conformations for transcription if the DNA is either overwound (gyrase)(42), or underwound (topoisomerase I) (43). An increase in the expression of both promoters simultaneously would therefore indicate topological stress in the chromosome consistent with the presence of persistent R-loops. We fused these promoters to a *lacZ* reporter gene and integrated these reporter cassettes into the *attB* site of the chromosomes of both wild-type and Δ*rnhC M. smegmatis* strains. Reporter activity from gyrase and topoisomerase promoters was increased in the Δ*rnhC* strain compared to wild-type, by 3-fold and 1.7-fold respectively (*p* < 0.05) (Figure 1B). Notably, these increases in activity were abolished in the complemented knockout strain, suggesting that chromosomal R-loops were indeed effectively processed by the complementing *rnhC* gene provided *in trans*.

### Loss of either *rnhA* or *rnhC* results in R-loop accumulation in *M. smegmatis*

To quantify R-loops in the RNase HI depletion strains, we isolated total nucleic acid from cultures of both Δ*rnhA* and Δ*rnhC* strains grown to mid-log phase and quantified the relative amounts of R-loops in each, using South-Western blots with a monoclonal antibody specific for DNA:RNA hybrids (44). The specificity of the antibody for RNA:DNA hybrids was confirmed by subjecting samples to RNase HI digestion prior to detection and noting the loss of signal (**Figure 2A**). R-loops were detected at low levels in the parental strain, and accumulated in the Δ*rnhA* mutant by a factor of ~2-fold, and in the Δ*rnhC* mutant by a factor of ~11-fold (**Figure 2B**). This indicated that both RnhA and RnhC act as part of a protective system to remove R-loops in replicating and actively transcribing cells. In both cases, the accumulation of R-loops also indicated that neither the remaining RNase HI isoform, nor other potential hybrid removal mechanisms such as the DNA helicase RecG (45), are able to compensate fully for the loss of either RNase HI enzyme.

**Figure 2:**
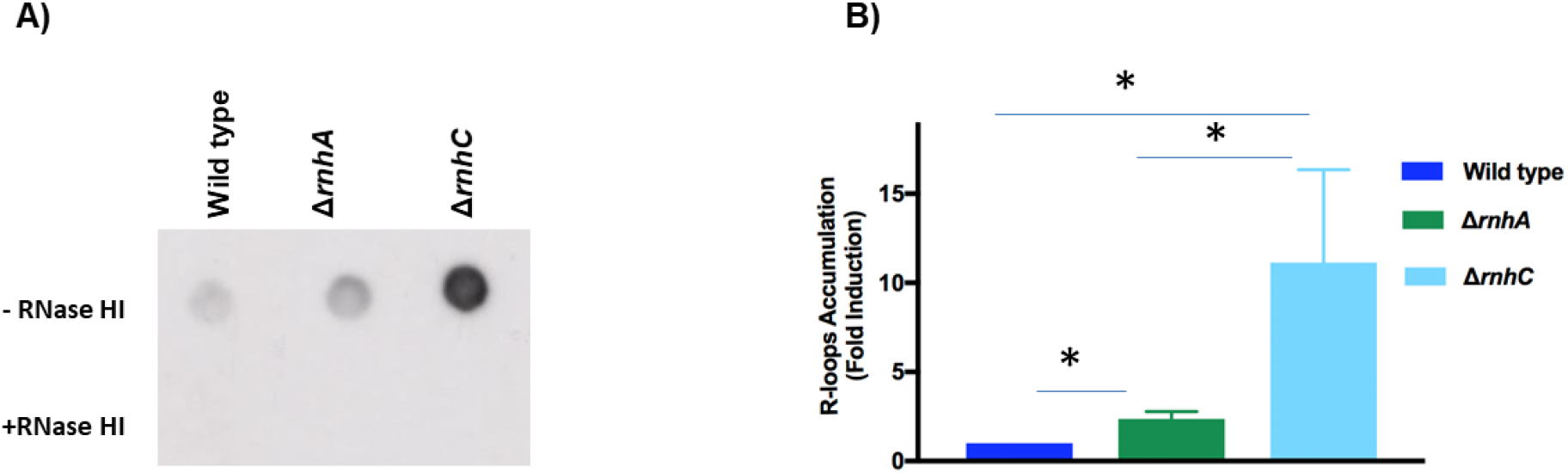
R-loop quantitation in wild type, Δ*rnhA* and Δ*rnhC* strains of *M. smegmatis* mc^2^155. **A.** Dot-blot analysis of R-loop accumulation: total nucleic acid from each strain was spotted onto the membrane and detected using a RNA:DNA hybrid-specific monoclonal antibody. Controls (+RNase HI) were treated with *E. coli* RNase HI before spotting. **B.**The amounts of R-loops in the wild type, Δ*rnhA*, and Δ*rnhC* strains were quantitated using Image Lab (BioRad). Relative amounts are shown normalized to the wild type. Data shown are representative of the average of three independent experiments with standard deviations indicated by error bars. The statistical significance of the differences are assessed using Student’s unpaired *t*-test calculated by GraphPad Prism 7. *, *P* < 0.05

### Plasmid replication exacerbates the R-loop burden in the *M. smegmatis* Δ*rnhC* strain

To confirm that loss of RNase HI activity was responsible for the observed increase in the number of R-loops in the Δ*rnhA* and Δ*rnhC* strains, we tested whether either *rnhA* or *M. tuberculosis rnhC* provided *in trans* could complement the Δ*rnhA* and the Δ*rnhC* mutants, respectively (**Figure S1 A/B**). As expected, complementation with the plasmid-borne *rnhA* gene reduced levels of R-loops in Δ*rnhA* to a wild-type level. In contrast, introduction of a plasmid carrying *rnhC* into the Δ*rnhC* strain conferred a 3-fold *increase* of R-loops over Δ*rnhC* without plasmid. A similar effect was obtained by transforming the Δ*rnhC* strain with the plasmid containing *rnhA,* or with vector containing the cobC domain only (**Figure S1C**). This suggested that the increase in R-loops was a function of plasmid carriage in Δ*rnhC*, and possibly unrelated to plasmid cargo.

We carried out whole genome sequencing of the *ΔrnhC* strain complemented with *rnhC* which determined that this strain was isogenic to *ΔrnhC*, except for the plasmid, and that the plasmid was present at between 24-48 copies per cell. Typically, pAL5000-based plasmids are present at 3-5 copies per cell (46), but a high copy number variant has been isolated. No mutations were found in the plasmid compared to the published sequence that could account for the increased copy number (47). This suggested that loss of RnhC activity affected plasmid copy number in *M. smegmatis* through a physical rather than genetic mechanism. Since the chromosomally-located reporters *gyrB::lacZ* and *topA::lacZ*, showed wild-type levels of topoisomerase activity in Δ*rnhC* p*rnhC*, but only report on the topology status of the chromosome, we concluded that the increased number of R-loops detected in the complemented *ΔrnhC* strain must therefore be plasmid-borne. It is possible that increased hyper-negative supercoiling of the plasmid, either through localised transcription effects (48) and/or titration of RNase HI activity to the chromosome (49), may have affected binding efficiencies of some of the proteins involved in plasmid replication, resulting in increased initiation frequencies, as well as promotion of R-loop formation.

Regardless of the precise mechanism, since the chromosomal DNA topology stress resulting from loss of *rnhC* could be fully complemented, we concluded that the plasmid copy number was well suited to the cellular requirements for RNase HI activity, and the supercoiling status of the plasmid might even reflect a useful cellular feedback mechanism to titrate the level of RNase HI appropriately. For these reasons, subsequent experiments were carried out using these constructs and strains. The contrasting behaviour of the complemented Δ*rnhA* and Δ*rnhC* strains also underscored the different physiological consequences of the loss of *rnhA versus rnhC* which may be a function of the severity of R-loop accumulation.

### Inhibition of transcription by low levels of rifampicin unexpectedly increases the accumulation of R-loops

Since transcription is a prerequisite for R-loop formation, we hypothesized that transcriptional inhibition would reduce the number of R-loops in the cell. Rifampicin inhibits transcription at the initiation step and does not affect RNA polymerase complexes that have added more than three ribonucleotides from proceeding to active transcription (50).

We exposed mid-log phase cultures of wild-type and Δ*rnhC* strains of *M. smegmatis* to rifampicin for 1 hour before harvesting the cells and quantitating the R-loops present in them. Rifampicin at a concentration equivalent to the MIC_90_ value for *M. smegmatis* (4.2 μM or 3.5 μg.ml^-1^) abolished R-loops, a result consistent with complete inhibition of transcription (**Figure 3**). Surprisingly, both the wild-type and Δ*rnhC* strains contained significantly increased amounts of R-loops when exposed to a concentration of rifampicin equivalent to the MIC_50_ value (1.2 μM or 1 μg.ml^-1^). A pronounced response to rifampicin exposure was shown by the Δ*rnhC* strain, with the number of R-loops increasing by ~3-fold compared to the non-treated Δ*rnhC* strain. Strikingly, even at a rifampicin concentration equivalent to MIC_25_ (0.6 μM or 0.5 μg.ml^-1^), more R-loops accumulated in Δ*rnhC* cells compared to untreated cells, indicating an overload of the cellular systems required for R-loop resolution. This also suggested that the low number of R-loops observed in wild-type cells at this rifampicin concentration was not due to lack of R-loop formation, but rather was reflective of efficient resolution by RNase HI. Notably, R-loop resolution is clearly impaired even in wild-type cells exposed to rifampicin at the MIC_50_. These results indicate that under these conditions, sub-lethal concentrations of rifampicin cause R-loop accumulation, a phenomenon that has not previously been described, and is opposite to the effect of a lethal concentration of rifampicin.

**Figure 3.**
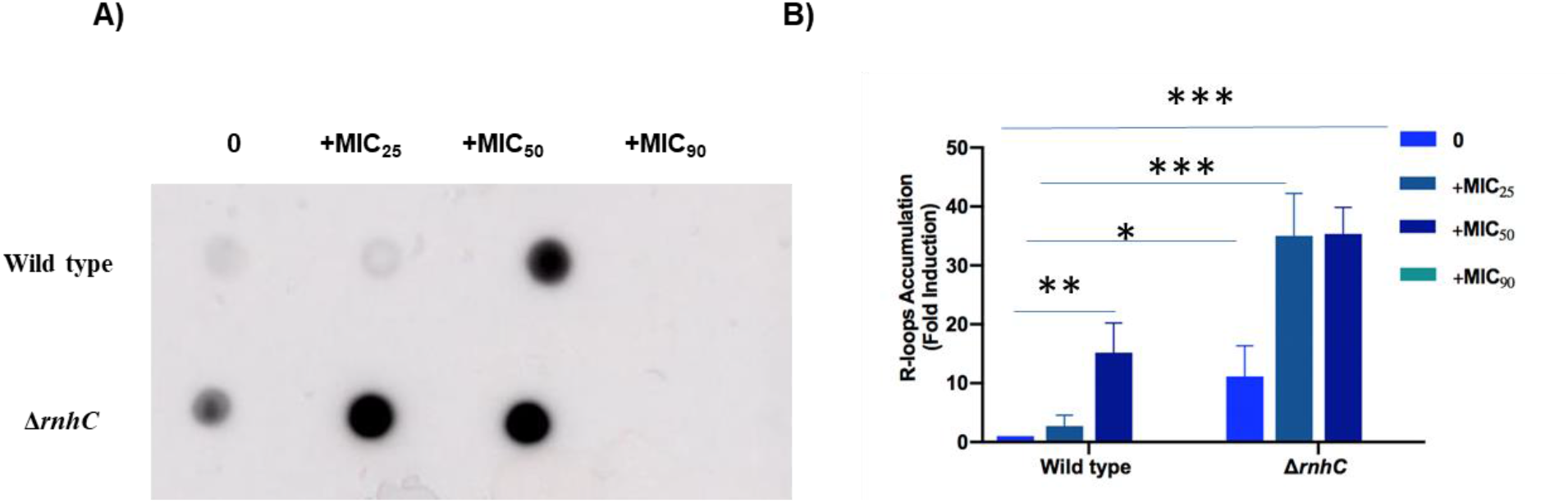
R-loop quantitation in wild-type and Δ*rnhC* strains of *M. smegmatis* mc^2^155 after exposure to various concentrations of rifampicin. **A.** Dot blot analysis of R-loop formation. **B.** Quantification of R-loops observed in the dot blot, normalized to the wild type strain in the absence of rifampicin. Data shown are representative of the average of three independent experiments with standard deviations indicated by error bars. The statistical significance of the differences assessed using Student’s unpaired *t*-test calculated by GraphPad Prism 7. \**, P* < 0.05; **, *P* < 0.01; ***, *P* < 0.001

### Loss of RNase HI is synergistic with transcriptional inhibition

To better understand the cellular consequences of rifampicin-induced R-loop accumulation, we carried out a dose-response assay to assess the extent to which RNase HI contributes to the survival of cells exposed to rifampicin. We found that wild-type *M. smegmatis* is inhibited as expected by previously established concentrations of rifampicin (51, 52). In contrast, loss of *rnhC* strongly sensitised the knock-out strain to rifampicin, reducing the MIC50 almost 100-fold, from 1.2 μM to 18 nM (Figure 4A). This sensitivity was fully complemented by the provision of *rnhC in trans*. To confirm that this remarkable loss of viability under rifampicin stress is due to the loss of the RNase HI domain rather than the CobC domain of RnhC, which is involved in vitamin B12 biosynthesis (20), the assay was repeated in the presence of vitamin B12 in the growth medium. Supplementation with vitamin B12 altered the MIC50 slightly. but not significantly, from 18nM to 30nM, (Figure 4A), indicating that the rifampicin sensitisation was due to the depletion of RNase HI function.

**Figure 4:**
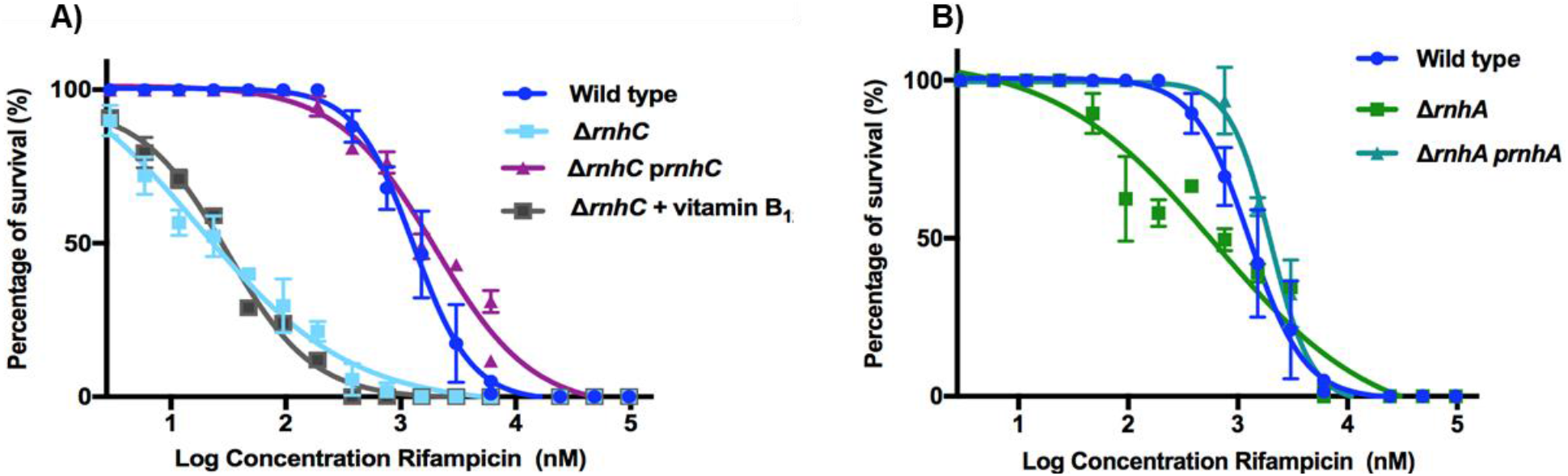
Dose-response curves of rifampicin killing in **A**) wild-type, Δ*rnhC* and **B**) Δ*rnhA* strains of *M. smegmatis* mc^2^155 and the respective complemented strains. Data shown are representative of the average of three independent experiments with standard deviations indicated by error bars. Vitamin B_12_ was included in the growth medium at 10 μg.ml^-1^.

As the Δ*rnhA* strain of *M. smegmatis* showed a smaller accumulation of R-loops than the Δ*rnhC* strain (**Figure 2**), we asked whether the loss of RnhA could also confer increased sensitivity to rifampicin. *M. smegmatis* Δ*rnhA* showed a ~3-fold increase in sensitivity to rifampicin compared to wild-type, which was fully complemented by the provision of *rnhA in trans* (**Figure 4B**). Together, these findings indicated that depletion of RNase HI activity was responsible for the increased sensitivity of *M. smegmatis* to transcriptional inhibition, and that loss of RnhA had a smaller contribution to this phenotype than the loss of RnhC. In fact, it was remarkable that such an apparently small contribution to R-loop metabolism still conferred synergy with rifampicin, underscoring the vulnerability of mycobacterial cells to even small amounts of RNase HI depletion.

### Loss of RNase HI also sensitizes *M. smegmatis* to moxifloxacin and streptomycin

R-loop formation can also be affected by uncoupling transcription and translation (53, 54), or by altering DNA supercoiling in the genome (5, 55). Therefore, we tested the sensitivity of the *M. smegmatis* Δ*rnhC* strain to streptomycin (an inhibitor of translation), amsacrine (a topoisomerase I inhibitor) and moxifloxacin (a DNA gyrase inhibitor) (**Figure 1A**). As a control, we investigated the sensitivity of the Δ*rnhC* strain to isoniazid, which disrupts mycobacterial cell wall synthesis by inhibiting mycolic acid synthesis (56).

The loss of the *rnhC* gene significantly reduced the MIC_50_ for streptomycin, from 31 nM in the wild-type to 17 nM in the Δ*rnhC* strain (*P* < 0.05), and to moxifloxacin, from 194 nM to 62 nM (**Figure 5A,B**). (*P* < 0.05). In contrast, the loss of *rnhC* did not result in a significant increase in the sensitivity of *M. smegmatis* to amsacrine (MIC_50_ = 160 μM for the wild-type and 140 μM for the Δ*rnhC* strain, *P* > 0.05) (**Figure 5C**). As expected, the loss of the *rnhC* gene did not significantly affect the MIC_50_ to isoniazid (85μM for wild-type and 63 μM for the knock-out strain, *P* > 0.05) (**Figure 5D**).

**Figure 5.**
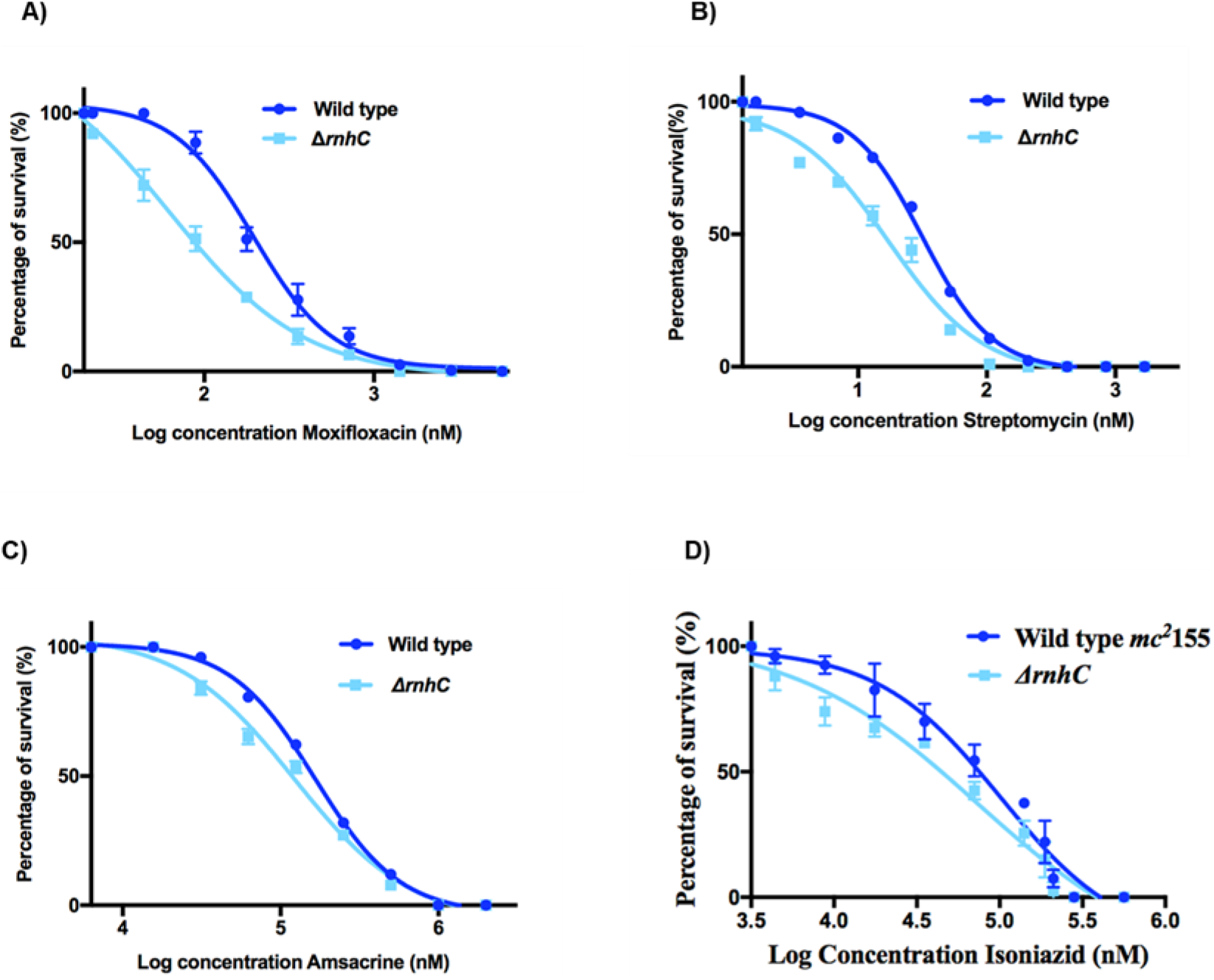
Dose-response curves of antibiotic killing for **A**) streptomycin, **B**) moxifloxacin,**C**) amsacrine and B) isoniazid in wild-type and ΔrnhC strains of *M. smegmatis* mc^2^ 155. Data shown are representative of the average of three independent experiments with standard deviations indicated by error bars.

Taken together, these results suggest that RNase HI inhibition would improve the efficacy of rifampicin, streptomycin and moxifloxacin in a synergistic manner. In all cases, the increase in efficacy arising from RNase HI depletion was more pronounced at sub-MIC concentrations of antibiotic, which is consistent with a requirement for some transcription to occur for the effect to be seen.

### Drug candidates with diverse scaffolds inhibit recombinant *M. tuberculosis* RNase HI

The enhanced antibiotic activity observed when RNase HI activity is depleted in *M. smegmatis* indicated that RNase HI inhibition is likely to be compatible with existing antibiotics used for *M. tuberculosis* therapy. Effective RNase HI inhibitors would be of particular interest as synergistic partners for rifampicin, which is a key first-line antibiotic used to treat tuberculosis. Rifampicin is a powerful inducer of hepatic cytochrome P450 enzymes, which leads to antagonistic interactions with many other drugs (57). Hence, any drug that sensitizes *M. tuberculosis* to rifampicin also has the potential to improve therapeutic outcomes by decreasing the dose of rifampicin required.

Recently, specific inhibition of HIV RNase HI was reported (14–16), preventing viral replication in cells, and supporting the possibility that small molecule compounds could be similarly discovered for bacterial RNases HI. The drug discovery effort against HIV RNase HI has identified promiscuous chemical scaffolds that not only have activity on HIV RNase HI *in vitro*, but also have activity on *E. coli* RNase HI or human RNase HI. We therefore selected a small library of 33 inhibitors of the RNase H activity of HIV-1 RT (**Table S3**) which we screened for activity against recombinant *M. tuberculosis* RNase HI. Ten compounds were found to inhibit *M. tuberculosis* RNase HI at a concentration less than 100μM (**Table S3**).

### Rifampicin synergy in whole-cell screens can eliminate off-target compounds

The ten most effective inhibitors of *M. tuberculosis* RNase HI *in vitro* were then tested for their antimicrobial activity against *M. tuberculosis* in a growth inhibition assay. All showed weak or no growth inhibition of *M. tuberculosis* when tested alone, which could be consistent with their modest inhibition of RNase HI *in vitro,* or with off-target activity. However, we hypothesised that on-target inhibition of *M. tuberculosis* RNase HI *in vivo* would phenocopy the genetic depletion of RNase HI observed in *M. smegmatis,* by increasing the sensitivity of *M. tuberculosis* to sub-MIC_50_ concentrations of rifampicin. Four compounds (NSC353720, NSC600285, NSC18806, and NSC99726) significantly potentiated killing of *M. tuberculosis* by sub MIC rifampicin indicating that they are likely to be inhibiting RNase HI *in vivo* (**Figure 6A and Table 1**). *In vitro* dose-response curves against *M. tuberculosis* RNase HI show that NSC353720 (IC_50_ = 14 μM) was the most potent inhibitor, while NSC99726 was the weakest inhibitor (IC_50_ = 45 μM) of this compound set (**Figure 6B**).

**Figure 6.**
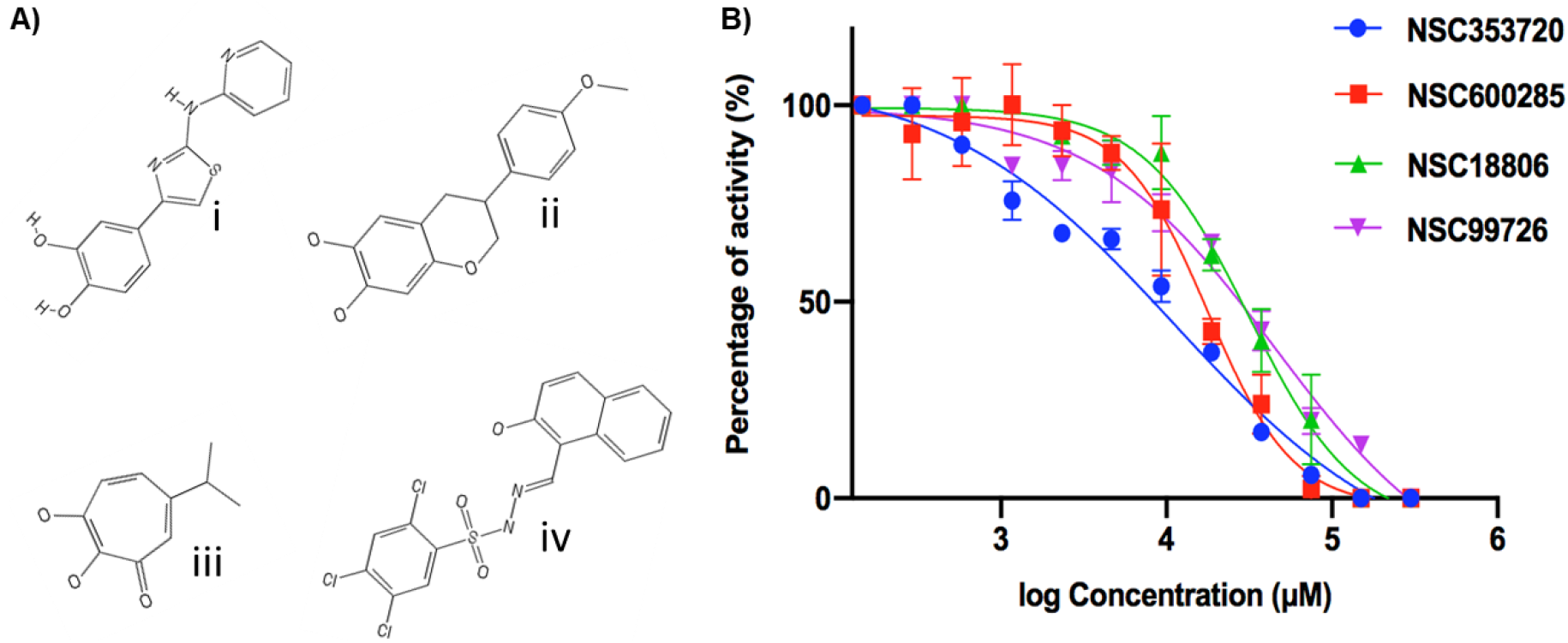
Chemical structures (**A**) and dose-response curves (**B**) for NSC353720 (i), NSC600285 (ii), NSC18806 (iii) and NSC99726 (iv) against recombinant *M. tuberculosis* RNase HI. Data shown are the average of two independent experiments with standard deviations indicated by error bars

**Table 1.**
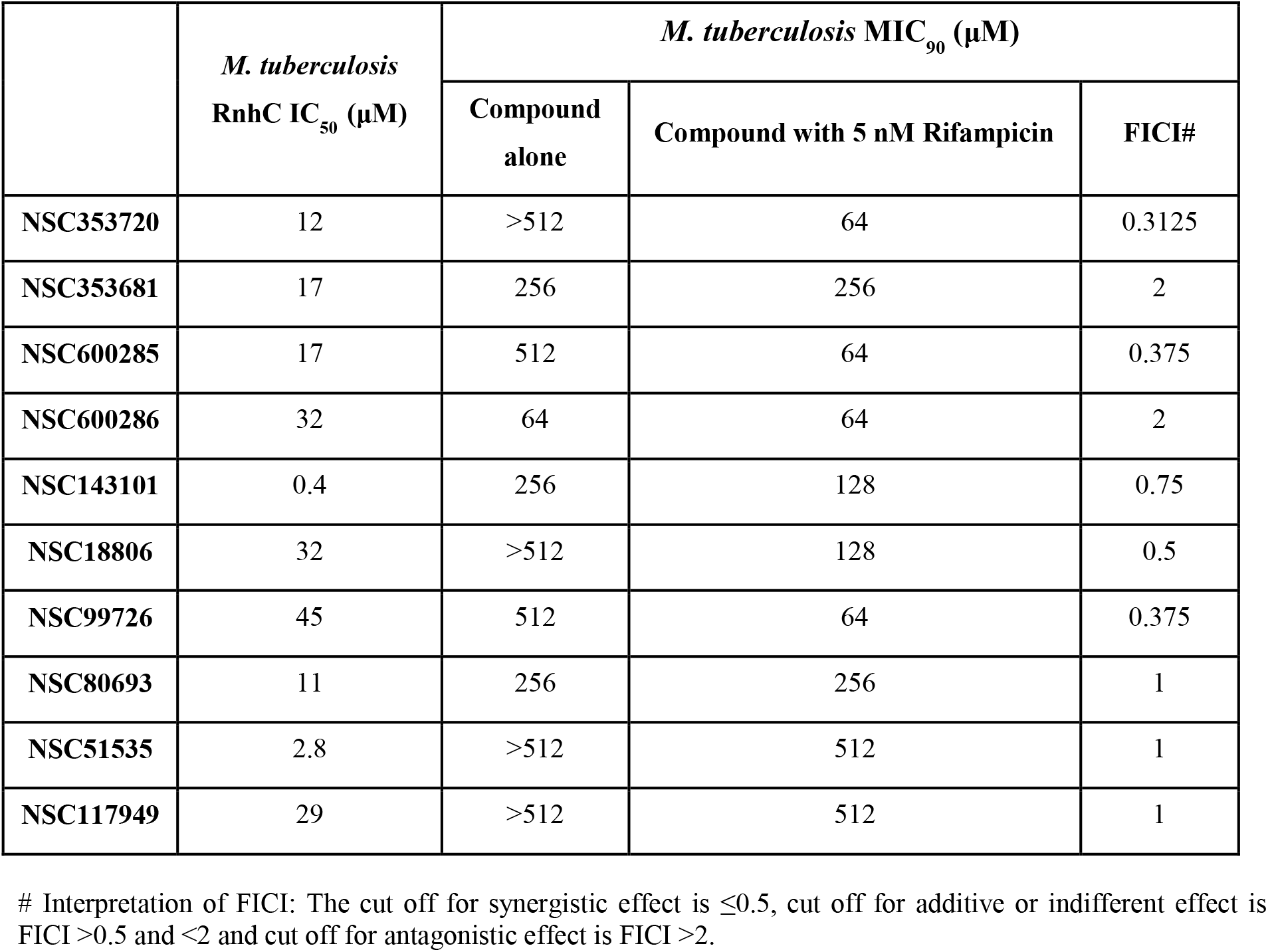
Inhibition of *M. tuberculosis* RNase HI recombinant protein and *M. tuberculosis* whole cells by *M. tuberculosis* RNase HI inhibitors and their interaction with rifampicin.

|  | *M. tuberculosis*<br>RnhC IC<sub>50</sub> (μM) | *M. tuberculosis* MIC<sub>90</sub> (μM) |  |  |
| --- | --- | --- | --- | --- |
|  |  | Compound alone | Compound with 5 nM Rifampicin | FICI# |
| NSC353720 | 12 | >512 | 64 | 0.3125 |
| NSC353681 | 17 | 256 | 256 | 2 |
| NSC600285 | 17 | 512 | 64 | 0.375 |
| NSC600286 | 32 | 64 | 64 | 2 |
| NSC143101 | 0.4 | 256 | 128 | 0.75 |
| NSC18806 | 32 | >512 | 128 | 0.5 |
| NSC99726 | 45 | 512 | 64 | 0.375 |
| NSC80693 | 11 | 256 | 256 | 1 |
| NSC51535 | 2.8 | >512 | 512 | 1 |
| NSC117949 | 29 | >512 | 512 | 1 |

## Discussion

R-loops are genotoxic stresses for all cells and the resolution of these structures is critical for cell survival. RNase HI is the primary enzyme that resolves R-loops, although cells have evolved multiple mechanisms to reduce their occurrence or to repair them. In *M. tuberculosis*, RNase HI activity is essential for the growth of the bacterium *in vitro*, which makes it a potential drug target. Nevertheless, not much is known about the cellular consequences of RNase HI inhibition in the mycobacteria. In this study, we showed for the first time that depletion of RNase HI activity in *M. smegmatis* caused an accumulation of R-loops. Loss of *rnhC* had a greater effect on R-loop accumulation than loss of *rnhA*.

The involvement of RnhC, and to a lesser extent RnhA, in the resolution of R-loops in *M. smegmatis* indicates that both enzymes are part of a protective system to remove the RNA/DNA hybrids that form spontaneously in the genome during transcription, before they induce DNA damage. The ability of *M. tuberculosis rnhC* to fully complement the Δ*rnhC* phenotype in *M. smegmatis* strongly suggests that RnhC carries out the equivalent role in both bacteria. Our data is consistent with previous reports that showed loss of RNase HI activity in *Bacillus subtilis* (58), *Saccharomyces cerevisiae* (59, 60) or human cells (61) resulted in the accumulation of R-loops. R-loop formation is favoured by high G+C content, G/C skew and is promoted by G4 quadruplex structures. Although sites susceptible to R-loop formation have not yet been mapped in the mycobacteria, over 10,000 sites for G4-quadruplexes in the *M. tuberculosis* genome have been predicted (62). The high (>65%) GC content of the mycobacterial genomes is likely to favour both the formation and the persistence of R-loops. If mycobacteria are more prone than other bacteria to R-loop formation due to their high GC content, this could account for the essential nature of RNase HI in the mycobacteria since R-loops promote replication-transcription collisions (6, 7, 9, 58, 63), DNA recombination (6, 59, 64–66), DSBs (9, 66–68), and gene silencing.

The striking finding that low levels of transcriptional inhibition promote R-loop accumulation gives new insight into R-loop metabolism in general. It is well established that polymerases which stall or backtrack during transcription are susceptible to forming R-loops (40, 41, 69–71), and that R-loops formation is able to stall a translocating polymerases. The “twin domain” model of transcription shows that DNA in the wake of RNA polymerase becomes negatively supercoiled (*i.e.* underwound) and that DNA ahead of it becomes positively supercoiled (*i.e.* overwound) (72–74). Hence, in actively transcribed genes which contain multiple polymerase complexes, the wake of unwound DNA from one polymerase is rewound by the following enzyme, thus reducing the need for topology-modifying enzymes and protecting this inter-polymerase region of DNA from R-loop formation (**Figure 7A**). Thus a topological balance exists in the DNA between actively transcribing RNA polymerases, which is disrupted if a polymerases is stalled.

**Figure 7.**
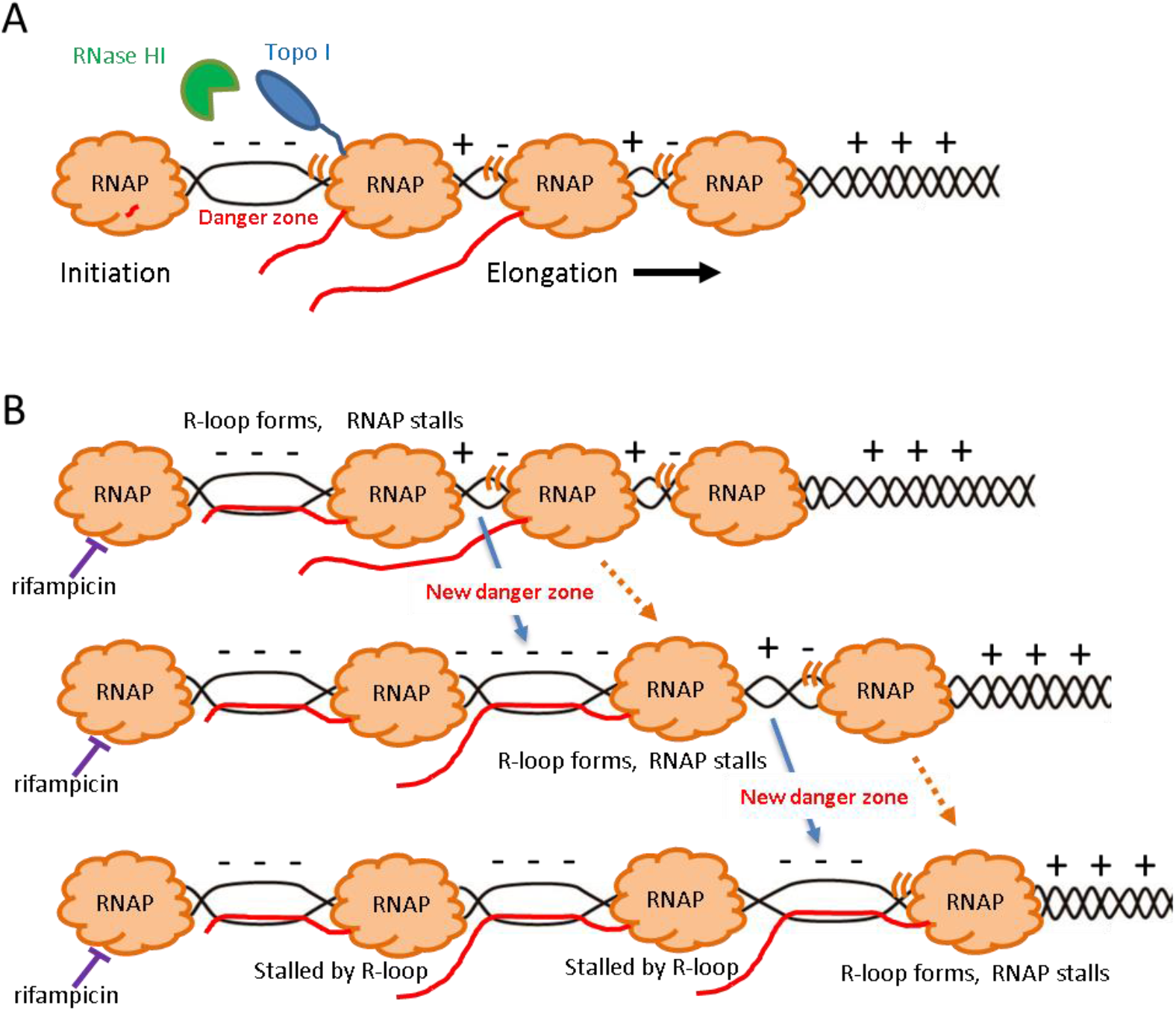
“Domino effect” model of R-loop formation under rifampicin stress. **A.** A negatively supercoiled DNA stretch (- - -) opens between a stationary RNAP (such as one in the promoter region, but can be paused anywhere) and a mobile RNAP (motion indicated by ‘((‘). This danger zone for R-loop formation is policed by topoisomerase I (blue, shown only once for clarity), which competes with mRNA (red line; only shown twice for clarity) for underwound DNA, and by RNase HI (green). Zones between mobile RNAP are more likely to be neutral (+ −). **B.**Under rifampicin stress, the promoter-bound RNAP cannot start elongating, the danger zone becomes critically underwound, an R-loop forms and stalls its cognate RNAP. A new danger zone opens up between this stalled RNAP and its adjacent mobile RNAP. A new R-loop forms, stalling its RNAP, and the cycle continues until all mobile RNAP molecules are stalled by R-loops. Ribosomes are not shown.

Our results indicate that stalled RNA polymerases in the promoter region promote R-loop formation. It is clear that an R-loop cannot arise directly from the rifampicin-bound polymerase, as no more than 3 nucleotides have been added to the transcript. We hypothesize that the last active polymerase will generate an increasing amount of underwound DNA, which promotes R-loop formation, which stalls this polymerase. A domino effect may thus be created over the length of the gene (**Figure 7B**). The magnitude of this effect depends on the initial stalling event to be close to the beginning of the gene. In contrast, stalling a polymerase near to the end of the gene is unlikely to promote cascading R-loop formation since trailing polymerases will merely stack up neatly behind it until the initial blockage is cleared. Halting all transcription evenly will also have a lesser impact on R-loop formation. It is possible that pause sites in promoter regions might facilitate optimal spacing between polymerase molecules to promote productive transcription by preventing R-loop production. This domino model of formation of R-loops under rifampicin stress is a general, biologically plausible model that nonetheless will be affected by the propensity of the DNA sequence to form R-loops, and the availability of the complementary RNA to bind to DNA.

Our observation that transcriptional inhibition leads to the accumulation of R-loops may also open a window into the poorly understood mechanism by which rifampicin causes cell death. Transcriptional inhibition has often been equated to translational inhibition in terms of cellular consequences, as cell death is expected to arise primarily from the lack of protein product (75) and not from lack of transcription *per se*. Transcriptomics is a widely used approach to gain insight into the mode of action of an antibiotic, but as rifampicin inhibits transcription globally, this approach has only been used on rifampicin-resistant strains (76) that may have altered and variable gene expression as a result. As one of the effects of R-loop formation is gene silencing, this may be more profound in some genes than others, and thus RNA sequencing together with R-loop mapping may give alternative insights. Our model suggests that an important aspect of rifampicin’s bactericidal activity may be the genotoxic stresses inflicted by R-loop formation.

We showed that even partial loss of RNase HI activity through the deletion of *rnhC* or *rnhA* synergised extraordinarily well with transcriptional inhibition by rifampicin at concentrations below the MIC_90_ (Fig. 4). This reveals an unappreciated role for RNase HI in promoting survival and possibly persistence under rifampicin stress. Rifampicin is one of the front line drugs for TB therapy (2), and a vital component in resolving bacterial persistence *in vivo* (77). Notably, exposure to sub-MIC concentrations of rifampicin is thought to be instrumental in the emergence of antibiotic resistance in clinical settings. Our results suggest that inhibiting both RNase HI and transcription might act against the emergence of drug resistance, even in sub-MIC concentrations of inhibitors of both RNase HI and RNA polymerase. It also raises the possibility that RNase HI inhibition might rescue rifampicin as a treatment option in rifampicin-resistant bacteria, by increasing their sensitivity to its cellular effects.

Partial loss of RNase HI activity also enhanced killing with moxifloxacin and streptomycin (Fig. 5A&B), and was compatible with isoniazid (Fig. 5D), indicating that in addition to providing a new target for drug development and enhancing first-line therapy, RNase HI inhibition would be a useful adjunct to second-line therapy as well. Uncoupling of transcription and translation is known to enhance R-loop production (54), and ribosomes can actively reverse stalled polymerases that might be more prone to R-loop formation (78). It is noteworthy however, that the combination of translational inhibition and RNase HI depletion does not affect the cell as potently as the combination of transcriptional inhibition and RNase HI depletion. We speculate that in the context of actively translocating RNA polymerases, the opportunity for R-loop formation due to translational inhibition is much reduced compared to the propensity for R-loop formation arising from stalled RNA polymerases, resulting in less synergy.

Many antibiotics, including rifampicin, are antagonistic with moxifloxacin (79) so the synergy of the latter with RNase HI depletion is both intriguing and valuable. We hypothesize that the synergy of moxifloxacin with RNase HI depletion might result from an increased number of gyrase/DNA complexes due to the 3-fold increased expression of DNA gyrase (Fig. 1B), resulting in a higher DSB burden (80).

Our finding that RNase HI depletion does not potentiate amsacrine lethality (Fig. 5C) contrasts with the synthetic lethal effect in *E. coli* of the loss of RNase HI and topoisomerase I (5). This apparent anomaly could be due to various factors. First, the expression of topoisomerase I is less effectively induced in the Δ*rnhC* strain than DNA gyrase I, which would be consistent with a topology-ameliorating effect of R-loops that has recently been proposed (81). Secondly, amsacrine is an uncompetitive inhibitor of topoisomerase I which binds to the topoisomerase I:DNA complex (82). Since topoisomerase I and mRNA are both localised by RNA polymerase, and are both competing for the same underwound DNA substrate, any R-loop formation would effectively remove underwound DNA as a substrate for topoisomerase I, and proportionally reduce the binding of amsacrine to a DNA:topoisomerase I complex, an opposite effect to that seen with moxifloxacin. A final possibility is that, although amsacrine is on-target in eukaryotic cells, and has been shown to inhibit *M. tuberculosis* topoisomerase I *in vitro* (82), it has not formally been shown that it does the same in *mycobacterial* cells. Topoisomerase I is an emerging drug target for development of anti-TB therapeutics. We predict that competitive inhibitors of topoisomerase I, which would more accurately mimic loss of the *topA* gene, might still be effective synergistic partners for RNase HI inhibitors.

The housekeeping nature of the enzymes involved in these synergistic combinations suggests that a wider application for combination therapeutics including RNase HI inhibitors also exists for other organisms. The loss of the *rnhC* gene in *Listeria monocytogenes* correlates with a loss of virulence in mice (58), and in *E. coli*, both streptomycin-resistant strains and rifampicin-resistant strains were competitively disadvantaged in Δ*rnhA* backgrounds (13). This indicates the potential for therapeutic aspects of RNase HI inhibition in bacteria even where RNase HI activity is not essential under laboratory conditions, or where rifampicin tolerance is high.

This study identified four HIV RNase HI inhibitors that have activity against *M. tuberculosis* RNase HI *in vitro* and show synergy with rifampicin in a whole cell assay (Fig. 6). The compound NSC600285 in particular is known to affect human RNase HI *in vitro* (bioassay ID: 5185715), but is not toxic to eukaryotic cells, unless the cells are deficient in DNA replication or repair, such as the Δ*rad50* and Δ*rad18* yeast strains (bioassay ID: 3254803). This supports an on-target effect in eukaryotic cells, but indicates that modifications are necessary to develop these chemical scaffolds for higher affinity and specificity for *M. tuberculosis* RNase HI. However, the inhibition we observe indicates that these scaffolds can penetrate both eukaryotic and prokaryotic cell walls, a key consideration for the delivery of antibiotics to intracellular pathogens such as *M. tuberculosis* and *L. monocytogenes*. In combination, these lines of evidence strongly support an on-target effect on RNase HI *in vivo* for these compounds, but the generation of resistant mutants to formally prove on-target *in vivo* activity will require the development of compounds with higher affinity. Crystal structures of the compounds in complex with *M. tuberculosis* RNase HI will allow more extensive structure-activity relationships to be established, and we are actively pursuing this avenue. The use of rifampicin as a sensitizing compound will be advantageous in high-throughput screening, both for isolating compounds that might otherwise inhibit RNase HI too weakly to be identified in a normal growth inhibition screen, and for prioritizing hits for further investigation and development.

As rifampicin is a naturally occurring antibiotic, produced by the actinobacterium *Amycolatopsis rifamycinica* (83), it is not likely to be present at lethal concentrations in its native environment. The model of rifampicin action proposed in this paper suggests that the maximum cellular impact on the target organism is achieved by stochastically halting transcription at its earliest point on the gene. Coupled with the physical protection of the initiation region from subsequent initiation events, this produces a formidable and subtle weapon since it uses the target cell’s transcriptional capacity to amplify the cytotoxic impact. Many further questions arise from this model. For example, does transcription arising from co-directional or divergent genes ameliorate or amplify this effect? Are terminal genes in operons disproportionally silenced? Does R-loop formation occur whenever there is a change in gene expression? Future work to answer these questions will help to further understand the intricacies of R-loop metabolism that this model implies.

In summary, this study validates RNase HI as a vulnerable, druggable target in the mycobacteria. It provides insight into R-loop metabolism in general, and specifically highlights the contribution that low-level transcriptional inhibition makes to R-loop formation. It also demonstrates the proof of principle that this is a novel cellular susceptibility, which can be utilised as an anti-bacterial strategy. Finally, it provides a starting point for the development of new anti-TB therapeutics based on re-purposing and expanding existing chemical scaffolds with favourable pharmaceutical properties.

## Acknowledgments and funding

We thank the Health Research Council of New Zealand (Grant 20/798), the Maurice Wilkins Centre for Molecular Biodiscovery, the South African National Research Foundation, the South African Medical Research Council, and the South African National Health Laboratory Service Research Trust for financial support. We thank Prof Deborah Williamson for helpful discussions.

