## Supplementary Tables and Figures for "RNase HI depletion strongly potentiates cell killing by rifampicin in mycobacteria"

### Supplementary information

Table S1 List of strains

| Strain | Genotype | Source |
| --- | --- | --- |
| E. coli DH5α | Cloning strain (the genotype: F^-^ Φ80*lac*ZΔM15 Δ(*lac*ZYA-*arg*F) U169 *rec*A1 *end*A1 *hsd*R17(r_k_^-^, m_k_^+^) *pho*A *sup*E44 *thi*-1 *gyr*A96 *rel*A1 λ^-^ | Invitrogen |
| E. coli BL21(DE3)pLysS | Protein expression strain  F-ompT hsdSB (rB-, mB-) galdcmrne131 (DE3) carrying pLysS (Cam^r^) | Invitrogen |
| *M. smegmatis* mc^2^155 | ept-1; high-frequency transformation mutant of M. smegmatis ATCC607 | (1) |
| *M. tuberculosis* mc^2^6230 | Δ*RD1* Δ*panCD* | (2) |
| *Ms* Δ*rnhA* | Derivative of mc^2^155 carrying inactivated allele of *rnhA* (*MSMEG_5562)* marked with Kan^r^ | This study . |
| *Ms* Δ*rnhC* | Derivative of mc^2^155 carrying inactivated allele of *rnhC* (*MSMEG_4305*) marked with Hyg^r^ | This study |
| Ms Δ*rnhA* p*rnhA* | Ms Δ*rnhA* complemented with p*rnhA*; Kanr, Hygr | This study |
| Ms Δ*rnhC* p*rnhC* | Ms ΔrnhC complemented with p*Rv2228c* | This study |
| *Ms* Δ*rnhC* pMV306G13+ LuxABCDE | *Ms* Δ*rnhC* containing pMV306G13+ LuxABCDE chromosomally integrated; Kan^r^ | This study |
| *M. smegmatis* mc^2^155  pMV306G13+ LuxABCDE | *M. smegmatis* mc^2^155 containing pMV306G13+ LuxABCDE chromosomally integrated; Kan^r^ | This study |

Table S2 List of plasmids

|  | Characteristic | Reference |
| --- | --- | --- |
| pMAL-C2 | Expression vector with IPTG inducible expression of the maltose binding protein as affinity tag and Factor Xa cleavage site upstream of the multiple cloning site; Amp^r^ | NEB |
| pAAZ-Rv2228c | pMAL-C2 plasmid modified by replacement of factor Xa site by His6-tag and 3C protease cleavage site; Rv2228c cloned in-frame for expression. | This study |
| pMV306G13+ LuxABCDE | Mycobacterial integrating vector containing the *luxABCDE* operon from *Photorhabdus luminescens*, Kan^r^ | (3) |
| pOLYG | pAL5000-based multicopy E. coli–Mycobacterium shuttle vector; Hyg^r^ | (4) |
| pEJ414 | Integrating plasmid containing promoterless *lacZ* gene, Km^r^ | (5) |
| p32ΔL-nuc | pOLYG with promoter and start codon of M. tuberculosis fbpA cloned as a 241 bp XbaI–BamHI fragment | (6) |
| p*rnhA* | pOLYG containing *rnhA* with 350 bp promoter region cloned as a *Bgl*II/*Hin*dIII fragment*.* | This study |
| p32GoriM | p32ΔL-nuc with 0.8 kb *aacC1* gene replacing *hyg* to give gentamicin resistance. | This study |
| p*Rv2228c* | Rv2228c cloned in-frame as a *Bam*HI/*Eco*R1 fragment to the p32ΔL promoter in p32GoriM | This study |
| pAAZ-*TopA* | pEJ414 containing 530bp upstream and 30bp coding sequence of *MSMEI_6157* fused in frame to *lacZ* | This study |
| pAAZ-*GyrBA* | pEJ414 containing 231bp upstream and 1000bp coding sequence of *MSMEI_0007* fused in frame to *lacZ* | This study |

Table S3. List of HIV RNase HI inhibitors and their activity against *M. tuberculosis* RNase HI at 100µM ( Active: compounds that show a reduction the activity *M. tuberculosis* RNase HI at 100µM, Inactive : no reduction in the activity was observed in the activity of M. tuberculosis at concentration 100µM of the compound )

| **compound** | **Activity against *M. tuberculosis* RNase HI** |
| --- | --- |
| NSC353720 | Active |
| NSC600285 | Active |
| NSC600286 | Active |
| NSC353681 | Active |
| NSC14543 | Inactive |
| NSC20410 | Inactive |
| NSC143101 | Active |
| NSC35676 | Inactive |
| NSC45382 | Inactive |
| NSC56351 | Inactive |
| NSC73300 | Inactive |
| NSC18806 | Active |
| NSC112200 | Inactive |
| NSC130796 | Inactive |
| NSC133457 | Inactive |
| NSC228148 | Inactive |
| NSC605657 | Inactive |
| NSC610984 | Inactive |
| NSC99726 | Active |
| NSC80693 | Active |
| NSC668394 | Inactive |
| NSC31892 | Inactive |
| NSC727447 | Inactive |
| NSC51535 | Active |
| NSC117949 | Active |
| NSC128437 | Inactive |
| NSC204474 | Inactive |
| NSC203867 | Inactive |
| NSC99722 | Inactive |
| NSC657715 | Inactive |
| NSC613575 | Inactive |
| NSC205497 | Inactive |
| NSC99727 | Inactive |

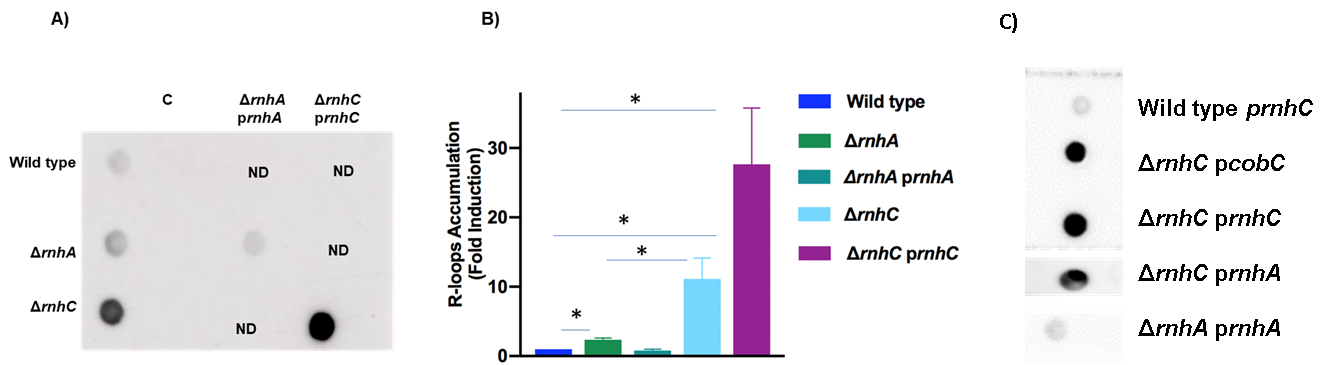

**Figure S1.** R-loop quantitation in wild type, ΔrnhA and ΔrnhC strains of M. smegmatis mc^2^155. **A.** Dot-blot analysis of R-loop accumulation: total nucleic acid from each strain was spotted onto the membrane and detected using a RNA:DNA hybrid-specific monoclonal antibody. Controls (lane C) were treated with E. coli RNase HI before spotting. **B.** The amounts of R-loops in the wild type, ΔrnhA, and ΔrnhC strains, and their respective complemented strains were quantitated using Image Lab (BioRad). Relative amounts are shown normalized to the wild type. **C.** R-loop accumulation in wild type, ΔrnhA and ΔrnhC strains of M. smegmatis mc^2^155 carrying alternative plasmids. ΔrnhC was transformed with gentamycin-based plasmids carrying either Mt rnhC (prnhC), or the cobC domain of Mt rnhC only (pCobC), or with a kanamycin-based plasmid carrying the rnhA gene cloned under control of its own promoter (prnhA). Data shown are representative of the average of three independent experiments with standard deviations indicated by error bars. The statistical significance of the differences are assessed using Student's unpaired t-test calculated by GraphPad Prism 7. *, P < 0.05
